# Extracellular vesicles-based point-of-care testing for the diagnosis and monitoring of Alzheimer’s disease

**DOI:** 10.1101/2024.03.31.587511

**Authors:** Xiang Li, Jie Chen, Yang Yang, Hongwei Cai, Zheng Ao, Yantao Xing, Kangle Li, Kaiyuan Yang, Abigail Wallace, James Friend, Luke P. Lee, Nian Wang, Feng Guo

## Abstract

Alzheimer’s disease (AD) is a debilitating condition that affects millions of people worldwide. One promising strategy for detecting and monitoring AD early on is using extracellular vesicles (EVs)-based point-of-care testing; however, diagnosing AD using EVs poses a challenge due to the low abundance of EV-biomarkers. Here, we present a fully integrated organic electrochemical transistor (OECT) that enables high accuracy, speed, and convenience in the detection of EVs from AD patients. We incorporated self-aligned acoustoelectric enhancement of EVs on a chip that rapidly propels, enriches, and specifically binds EVs to the OECT detection area. With our enhancement of pre-concentration, we increased the sensitivity to a limit of detection of 500 EV particles/μL and reduced the required detection time to just two minutes. We also tested the sensor on an AD mouse model to monitor AD progression, examined mouse Aβ EVs at different time courses, and compared them with intraneuronal Aβ cumulation using MRI. This innovative technology has the potential to diagnose Alzheimer’s and other neurodegenerative diseases accurately and quickly, enabling monitoring of disease progression and treatment response.

---

Alzheimer’s disease (AD) is a chronic and progressive neurodegenerative disease representing the main cause of dementia, and is quickly becoming one of the most expensive, lethal, and burdensome diseases of this century ^1^. The progression of AD is associated with amyloid plaques and tangles in the brain, which are composed of amyloid β (Aβ) and tau, respectively. These molecules are therefore considered promising biomarkers for AD diagnosis. However, limited by the blood-brain barrier (BBB), the acquisition of samples from the brain microenvironment often requires invasive techniques, such as lumbar puncture, that pose safety issues and incur patient suffering. Therefore, the diagnosis of AD currently relies on image-based detection methods, such as magnetic resonance imaging (MRI). Although this strategy is safe, convenient, and non-invasive, MRI is expensive and lacks the sensitivity to easily diagnose AD in its early stages or to monitor its progress. When detectable imaging changes occur in the brain, the disease has usually progressed beyond the best time for medical intervention.

How to find a carrier that can freely cross the BBB and carry disease biomarkers becomes the core issue to realize early diagnosis and process monitoring of AD. Fortunately, Extracellular vesicles (EVs) have been shown to have such potential, which are nano/microscale membrane-enclosed structures secreted by cells into the extracellular space, widely distributed in various biofluids, including serum, urine, cerebrospinal fluid, and milk^2^. EVs have been shown to play an important role as vehicles supporting long-range intercellular communication in the body, and are consequently recognized as promising circulating disease biomarkers, including neurodegenerative disease ^3-5^, traumatic brain injury ^6^, and cancer ^7-9^. EVs respond to pathological changes in the brain more accurately than plasma cytokines because of their ability to cross the blood-brain barrier ^10-12^, of particular interest in diagnosing brain-related diseases. Therefore, the construction of non-invasive, accurate, and simple methods based on EVs for point-of-care testing (POCT) may be crucial in the early detection and monitoring of brain-related diseases.

Despite the clinical potential of EVs, most conventional analytical methods lack sufficient sensitivity or specificity to be practical in the clinical use of EVs, principally because the EVs are so small (typically 50∼200 nm in diameter), rare, and highly heterogeneous. Especially in POCT applications, the requirements for sensing are particularly stringent, demanding ease of operation, specific loading volumes, rapid detection times, and high sensitivity. Especially facing to complex biological samples without pre-treated, uncontrolled and heterogeneous impurities in the background can harshly challenge the specificity of the sensor. Moreover, even a low percentage of false negatives and false positives can generate a large number of misdiagnoses among mass screening, which is a common scenario for POCT, leading to public panic. The classical approach is to locally enhance the target material in the sensing region via an exogenous field or guiding microstructures ^13-16^. Any technology that could potentially address the “bench to bedside” challenge of EV analysis needs to meet several conditions simultaneously: 1) speed, as it should have the ability to enrich bioparticles at the submicron level in minutes; 2) ease of use, so that untrained patients can correctly operate it after reading the instructions; 3) compatibility, as it must have the ability to work with untreated or simple pretreated samples; and 4) low cost, an important basis for rapid adoption and broad use.

Organic electrochemical transistors (OECT) are a combination of high-performance transducer and amplifier that convert biological signals into electrical signals, and have shown excellent potential in POCT applications ^17, 18^. A typical OECT is comprised of three terminals: the source electrode (SE), the drain electrode (DE), and the gate electrode (GE). A conductive organic material is introduced to connect the source and drain, forming the organic channel, and an electrolyte medium is used to connect the gate electrode and the organic channel. Compared with other types of thin-film transistors, the presence of the corresponding electrolyte double layer (EDL) at the gate-electrolyte-channel interface can provide a large gate-channel capacitance, and the penetration of the ions into the organic channel enables the OECTs to operate at a relatively low gate voltage but with good signal amplification ^19-21^. Biofunctionalization at either the organic channel or the gate electrode has been widely used for bioelectronic applications with good biocompatibility and specificity ^22-25^. In the classical OECT-based sensing system, the target bioparticles rely on diffusion and convective transport to bind to recognition sites on the surface. Thus, effective detection depends on a suitable initial concentration. The binding process becomes difficult when the concentration is too low, and the detection will then fail. Therefore, creating a high-concentration zone near the detection area is the key to achieving EV detection. To accomplish this aim, we consider acoustofluidics. Acoustofluidics is an interdisciplinary technology, driving matter or fluidics through acoustic pressure or acoustic streaming, which are well matched to the requirements in the pre-concentration of EVs, providing versatility, non-contact, biocompatibility, flexibility, low cost, and integrability ^14,26-32^. In recent years acoustofluidics has demonstrated extraordinary performance in pre-enrichment in highly constrained environments (sessile droplet and microfluidics).^33^ Moreover, the introduction of focused acoustic wave structures has pushed manipulation capabilities to the nanometer level.^34^

Here, we describe an integrated OECT-based sensor that enables high accuracy, speed, and convenience in detecting EVs from AD patients. We developed a self-aligned acoustoelectric enhancement system combining acoustics and electric fields to achieve the pre-concentration of EVs expressing specific proteins, thus enhancing the sensitivity and reducing the detection time. We tested tau-EVs from the serum of AD, showing the extraordinary performance of this integrated sensor in AD diagnosis with high speed, selectivity, and sensitivity. Further, we demonstrate the evaluation of AD disease progression based on this device in an AD mouse model (5xFAD), showing an age-dependent Aβ-EV increase in the collected serum profile, which was strongly correlated with the plaque loading data generated from high-resolution MRI. This innovative technology has the potential to diagnose AD and other neurodegenerative diseases accurately and quickly, enabling monitoring of disease progression and treatment response.

## Results

### Overview of integrated OECT sensor

The workflow and schematics of integrated OECT sensor are shown in **Fig. 1a-b**. The workflow can be divided into three parts: sample loading, enrichment, and detection. Briefly, 5 μL diluted serum obtained from AD patients is loaded into the reservoir on the chip, which contains an integrated focusing surface acoustic wave (FSAW) device and a specially designed OECT electrode, as shown in **Fig. 1b**. Then, EVs in the serum rapidly migrate toward the sensing region by the acoustic and electric fields. EVs carrying target proteins are captured by antibodies pre-modified in the detection region. Finally, their number is ultimately reflected in the results as a change in the shift of OECT current, which can be used as a basis for diagnosing AD in early stage; and multiple tests over time can reflect the progression of the AD disease. The core component of the system is the acoustic-electric enrichment module, as shown in **Fig. 1b**. The acoustic enrichment module is a pair of symmetrically focused IDTs (37.4 MHz) that generates a focused acoustic pressure at the focal point to concentrate EVs. In contrast, the electrical enrichment is realized by multiplexing the OECT electrodes. It is worth emphasizing that the OECT here was specially designed. Compared to classical OECT with two individual electrodes as the source and drain electrodes, the OECT in the integrated OECT sensor system have parallel forked-finger electrodes that cover the focused acoustic region to obtain the best performance. The simulation results of the distribution of the acoustic/electric field are shown in **Fig. 1c**. FSAW are localized fields with high intensity, while electric fields are fields of large scope with relatively weak intensity. They complement each other in terms of spatial distribution and intensity. In classical sensing systems, the EVs are randomly distributed among the sessile droplets and rely on Brownian motion-driven diffusion to contact the interface to react with the antibodies, which is inefficient and slow. In the integrated sensing system, we introduced an acoustic field and pulsed electric field into the detection system to achieve rapid enrichment of EVs, improve detection sensitivity, and shorten the detection time of OECT. Under the action of both acoustics and electric fields, the EVs in the sessile droplet are focused onto the surface of the detection area, forming the pattern of a parallel array of lines. As a result, the target EVs can efficiently and rapidly bind to the interfacial antibody to achieve sensing enhancement. We characterized the distribution of EVs in the sensing region by scanning electron microscopy (SEM) with and without acoustoelectrical enhancement (**Fig. 1d**). It can be seen that with the enrichment, the EVs are massively aggregated and arranged in bands. In contrast, EVs are sparsely and randomly distributed on the substrate without the acoustoelectrical field. We noticed that the heavily enriched EVs in the sensing region maintained an excellent morphology and an intact membrane, indicating that our system has both strong enrichment capability and good biocompatibility.

**Fig. 1.**
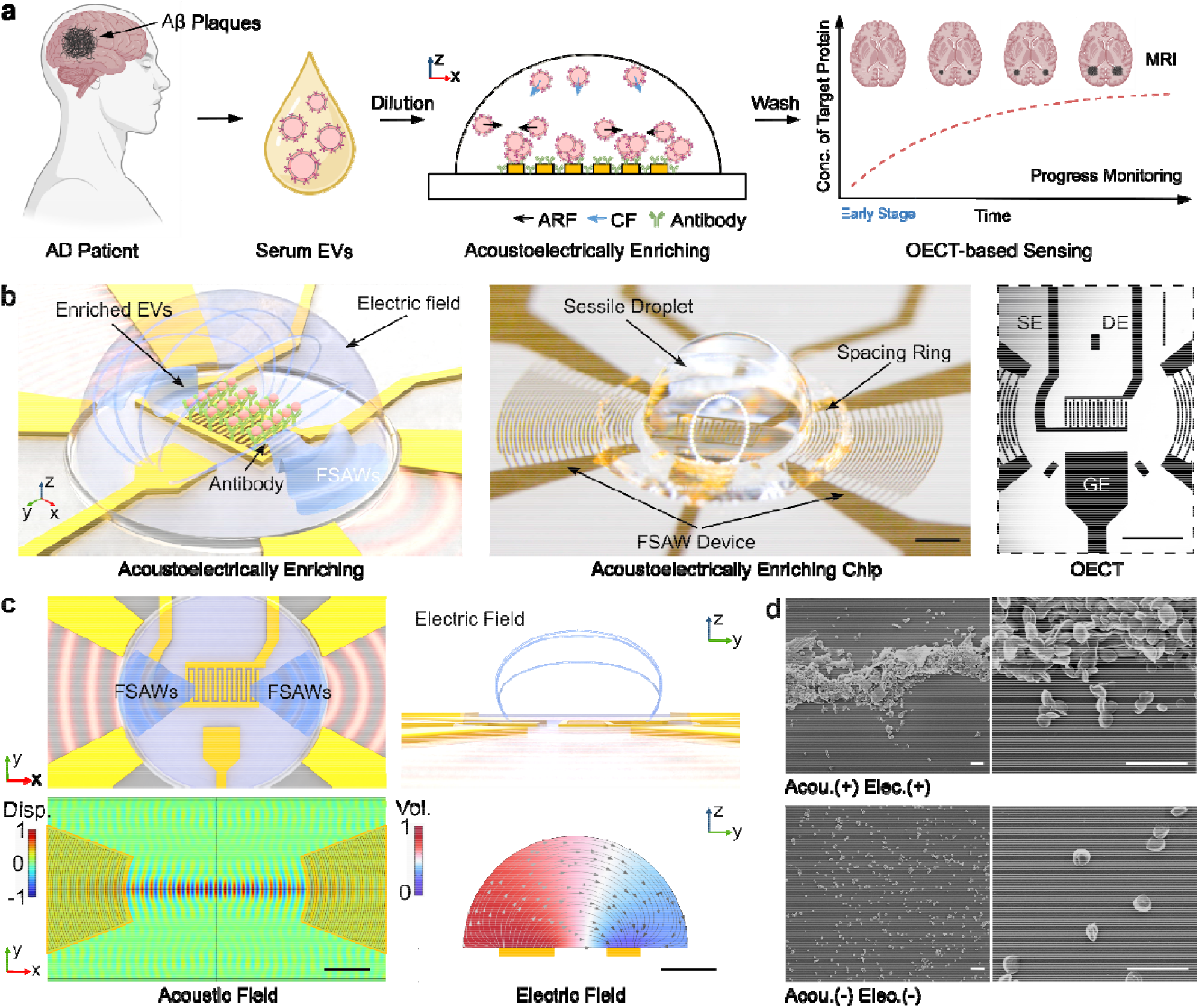
Point of care testing of serum extracellular vesicles (EV) in Alzheimer’s disease (AD). (**a**) Workflow of the integrated OECT sensor for diagnosis and monitoring of AD. (**b**) Schematics of EV enrichment through an acoustoelectrical manner. The focused surface acoustic waves (FSAWs) and the electric pulses rapidly enrich EVs for specific attachment onto the OECT coated with Aβ or Tau antibodies. The middle picture is the photo of the integrated OECT device and the detailed image in left shows the OECT sensor. The source, drain, and gate electrode are represented by SE, DE, and GE, respectively. Scale bar, 1 mm. (**c**) The simulation results of the distribution of acoustic/electric field. Scale bar, 500 μm. (**d**) Comparison of acoustoelectrically enhanced EV binding (acoustics on (Acou.(+)) and electrics on (Elec.(+))), with conventional EV binding (acoustics off (Acou.(-)) and electrics off (Elec.(-))). The SEM images demonstrate the distribution status of EVs in both binding modes.

### Characterization of acoustoelectrical enrichment

When the focused IDT is stimulated by an RF signal, two sets of traveling SAWs are each produced as a Rayleigh wave. The exponential decay of the amplitude with the depth of the substrate allowed the wave to confine most of its energy to the surface at the selected frequency in our design ^35^. The two focused SAW waves meet at the focal point to form a standing SAW, producing surface displacement from the SAW propagating along the x-axis that gradually intensifies toward the center of symmetry (**Fig. 1c**). The strongest vibration region is the focal point of the IDT, which is also the location of the OECT sensor. To visualize the acoustic field induced by the fabricated focused IDT device, we used 2 μm fluorescent polystyrene microspheres as tracer particles. The enrichment process is shown in Supplementary Fig. S1, where the fluorescent particles were enriched within ten seconds. The symmetrical axis of Supplementary Fig. S1b is regarded as the zero position of the x-axis. The Supplementary Fig. S1c curves indicate the simulated displacement (Si-Disp) and fluorescence intensity (FI) along the x-axis, respectively. The data was extracted from Supplementary Fig. 2b. The pattern of the enriched particles was strongly consistent with the displacement in the simulation results, indicating that we are indeed generating powerful FSAWs at the focal point and the particles were indeed enriched at the nodal positions.

To profile the enrichment process, it is important to understand the mechanism and characterize the performance of the hybrid acoustic-electric field. Here, we characterized and discussed the enrichment effect of acoustic and electrical enrichment separately using fluorescence-labeled EVs, which are purified from an AD patient’s serum by an EV purification kit (**Fig. 2a**). The size distribution and concentration of purified EVs were measured by nanoparticle tracking analysis (NTA) and dynamic light scattering (DLS), shown in Supplementary Fig. S2, which indicated that the purified EVs possessed the expected size properties. The fluorescence-label EVs were enriched with or without acoustics (Acou. -/+) and with or without electric fields (Elec. -/+) over 2 min. In pure acoustic enrichment (blue rectangle), the focused acoustic field triggers standing waves at the sensing location. The particles move toward the acoustic node under the action of acoustic radiation forces, thus achieving enrichment. In pure electrical enrichment (green rectangle), the EVs are attracted by the electric field toward the sensing region. In acoustoelectrical enrichment (red rectangle), which has both acoustic and electric fields, EVs are enriched to the surface of SE and DE under the dual action of the pulsed voltage and acoustic fields and specifically captured by the pre-modified antibody. The pattern of enriched EVs under the action of acoustics was highly consistent with the simulation results in **Fig. 1c**. Further, we extracted the FI of the fluorescent images along the x-axis to evaluate the enrichment performance (**Fig. 2b**). A white dashed rectangle represented the region of interest (ROI). The results showed that electric and acoustic enrichment have entirely different effects: The electric enrichment based on pulsed voltage is a homogeneous enrichment of vesicles on the sensing interface, reflected as a uniform enhancement of fluorescence; the acoustic enrichment is inhomogeneous, whereas EVs were patterned as parallel. Moreover, noticeable fluorescence strip enhancement is observed when the acoustic and pulsed electric fields are turned on simultaneously. In contrast, the fluorescence in the background increased, meaning the enrichment effect could be superimposed.

**Fig. 2.**
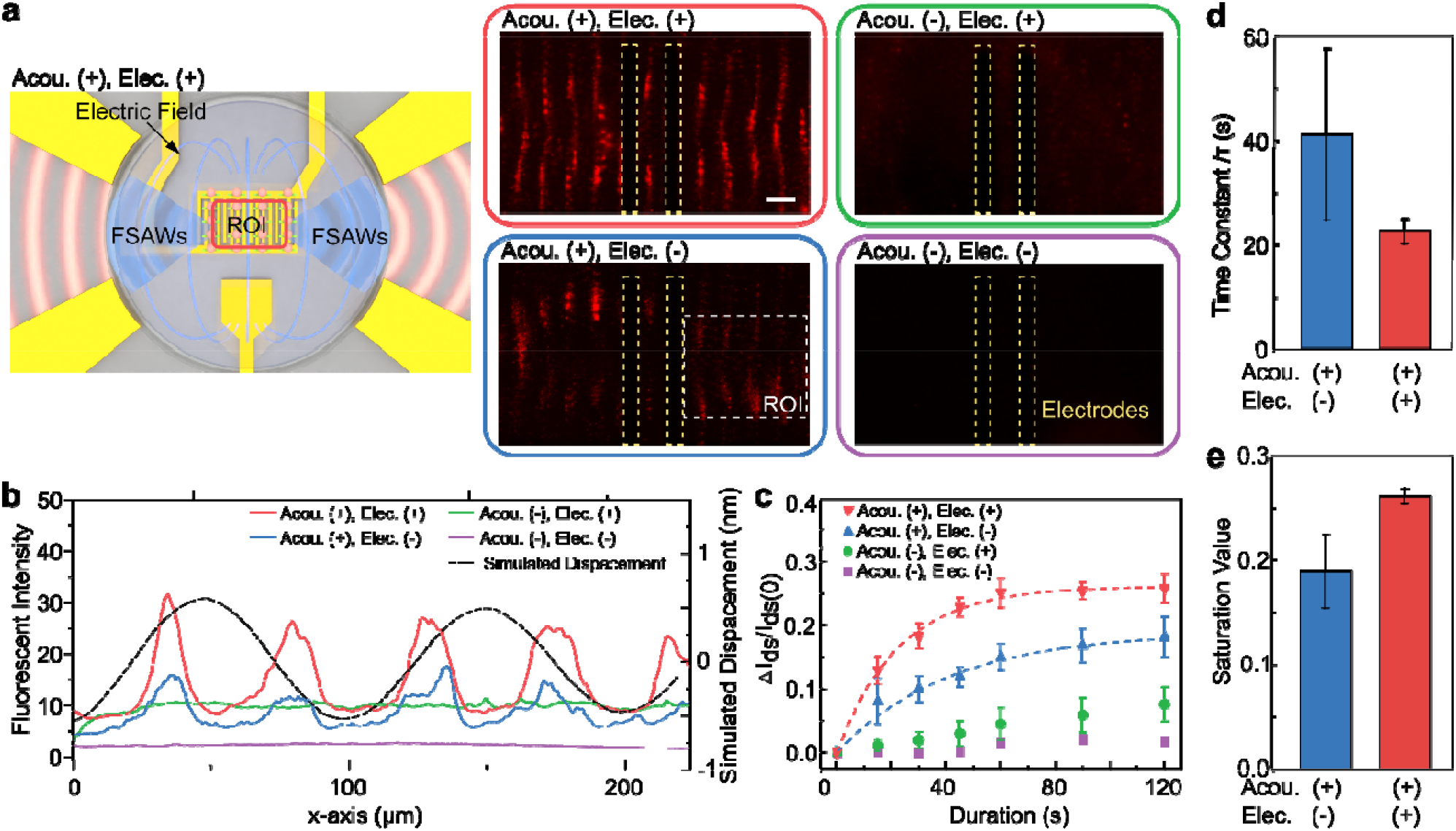
Integrated OECT sensors based on acoustoelectrical enhancement. (**a**) Fluorescence results of fluorescence-labeled EVs in the presence of acoustoelectrical enhancement. Acou(+) and Acou(-) indicate with/without acoustics, while Elec.(+) and Elec.(-) indicate with/without voltage pulse. The region of interest (ROI) is highlighted by a red rectangle in the cartoon image. The scalebar is 50 μm. (**b**) Fluorescent intensity extracted from the ROI (white rectangle in Fig. 2a) and corresponding simulated displacement of acoustic vibrations. The yellow rectangles represent the electrodes of OFET. (**c**) Relative change of the electric current between SE and DE of OECT as a function of incubation time under different conditions. The dashed curves indicate the response curves fitted using the exponential function. The time constant (**d**) and saturation value (**e**) of the fitting curves in (**c**).

Further, we tested the enhanced performance of this acoustoelectrical manner on the OECT-based sensing to evaluate the role played by acoustics/electrics. Hence, we tried to build the relationship between the normalized current shift of the transfer curve (Δ*I*_*ds*_*/I*_*ds*_*(0)*) and the incubation time. The transfer curve of the processed OECT sensor and the shift of transfer curves for purified EV detection are shown in Supplementary Fig. S3a and b, which showed a linear response over the operating voltage range. The purified EVs with a concentration of 4.6 × 10^7^/mL were dripped on the sensing area for a certain period (2 min) for incubation with/without acoustic field (frequency: 37.4 MHz, amplitude: 17 *V*_*pp*_) and with/without voltage pulse (frequency, 10 kHz; voltage, -0.5 V; pulse width, ten μs), and then, the transfer curve of the OECT was measured in PBS buffer after washing three times. This process on the same device was repeated several times with different incubation times. Based on these, the device performance as a function of accumulated incubation time was obtained, as shown in **Fig. 2c**. During the two-minute incubation time, in the group without both acoustics and pulsed electric fields, little change was seen in the conversion curve, indicating that the classical OECT sensor was incapable of rapid detection at this EV concentration. A tiny response could be seen when the pulsed electric field was turned on alone. When the acoustic field was turned on alone, a rapid rise in the curve was seen, and saturation was reached within two minutes of incubation, demonstrating the excellent performance of the focused acoustic field in enriching EVs. It is worth emphasizing that the two effects were superimposed when the electric and acoustic fields were turned on simultaneously. The electric field further promoted the enrichment performance of the acoustic field, shortening the saturation time and increasing the saturation value. The relationship can be fitted with an exponential function 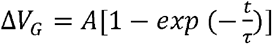, where *τ* is the time constant, which represents the time required to rise to a saturation value of 63.2%, and *A* is the saturation value of ΔI_ds_/I_ds(0)_. The *τ* of the “Acou.(+) Elec.(+)” group and the “Acou.(+) Volt.(-)” group is 22.4 s and 41.2 s, which indicates that the introduction of the pulsed voltage field can reduce the incubation time of the acoustic field by about 50%. Moreover, the saturation value also increased from 18.94% to 26.13%, which indicates that the introduction of the pulsed electric field amplifies the gain multiplier, improving the limit of detection (LOD). It is worth stating that the system can complete saturation adsorption of EVs in less than 2 minutes, which makes the device well-suited for rapid readouts in POCT, where it is difficult to require the user to precisely control the incubation time but a quantitative test is needed.

### AD diagnostics in serum samples

After evaluating the performance of the sensing enhancement, we further validated its sensing performance in clinical samples. We tested different concentrations of purified EV samples and serum from AD patients with a 2-minute incubation time to calculate the LOD of the integrated OECT sensing system (**Fig. 3a and b**), showing an excellent LOD (calculated as 3 x standard deviation of the baseline (PBS)) of 3.17 × 10^3^ /mL and 0.53 × 10^3^ /mL in PBS and serum, respectively. The serum testing group showed a “theoretically” better LOD due to its better linearity within the range of testing. However, the serum group reduced dynamic range as the I_ds_ shift was less than the purified EV group at higher concentration, as well as increased cut-off range due to the increased I_ds_ shift in the healthy control (the normalized I_ds_ shift was 0.050 ± 0.025 in the healthy serum group, while 0.015 ± 0.004 in the healthy purified EV group, **Fig. 3c**). As a result, the corresponding limit of quantification (LOQ) for direct serum EV detection was estimated to be 2.028 × 10^5^ /mL. This performance difference reflected the increase of possible interference within serum samples, such as plasma proteins and nano-sized lipids, that might have contributed to the non-specific binding of our current detecting mechanism (i.e., the affinity binding between antigen and antibody). Instead of relying on off-the-shelf antibodies, the selectivity of our system can be further optimized by incorporating more advanced biofunctionalization approaches, such as self-assembled monolayers and aptamers ^18, 36^.

**Fig. 3.**
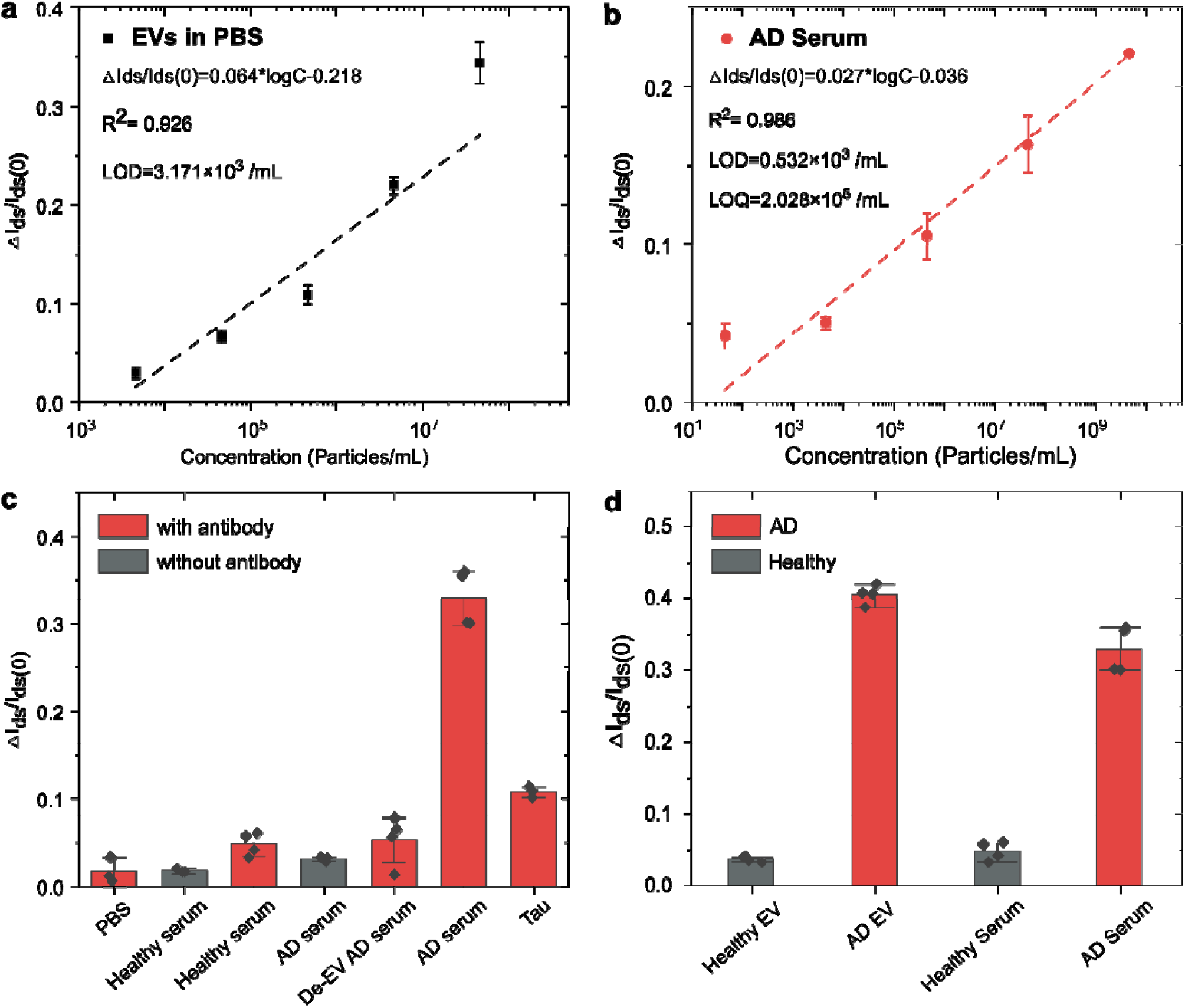
Detection of serum EVs from AD patients. (**a**)/(**b**) Sensitivity of POCT sensors in detecting tau EVs from purified EV sample/ AD serum. The points are means based on three independent replicates; bars represent s.d.. LOD, the limit of detection. (**c**) Specificity of the integrated OECT sensor. The testing results for AD serum from AD patients, health donors, EV-depleted serum, and PBS buffer. (**d**) AD diagnosis based on serum from patients. The sensing results for purified EVs and serum from health and AD patient donors.

To harness the high sensitivity of our system and reduce the impact of the individual patient difference, we directly diluted a minimal volume of serum samples that can be collected from the fingertip. We tested the serum response from AD patients at different dilutions with/without modified antibodies to find the optimal dilutions (Supplementary Fig. 4a). We defined the response of the unmodified device as the noise induced by non-specific adsorption and the response of the modified device as the signal. Thus, the signal-to-noise ratio (SNR) is the response of the modified device divided by the reaction of the unmodified device. The SNR at different dilutions is shown in Supplementary Fig. S4b. Considering the response value and SNR, we chose the 10^12^ times as the sample pretreatment parameter for the subsequent experiments.

To evaluate the specificity of this integrated sensor, serum samples from AD patients (n=3) and healthy individuals (n=3) were divided into two parts, one for EV purification by a commercial kit to obtain EVs in a PBS buffer before detection and the other for detection directly after dilution. The incubation time was 2 minutes. The relative change of normalized *I*_*ds*_ shift was demonstrated in **Fig. 3c**. There were two variables: whether or not the surface of the device was modified with an antibody (Positive: with antibody, Negative: without antibody) and whether or not the sample contained EVs from AD patients (Positive: AD serum, Negative: Healthy serum and De-EV AD serum), where De-EV AD serum indicates AD patient serum after removal of EVs. The results showed a significant response from the group with modified antibodies and EVs in the sample, indicating that this acoustoelectrically sensing-enhanced technique possesses excellent specificity.

Acoustoelectrical enhancement was further applied to diagnosing clinical samples, as shown in **Fig. 3d**. The relative changes in the transfer curve in the patient samples were much greater than that in healthy individuals. The results of purified EVs were consistent with those of diluted clinical samples, which indicated that the integrated sensor system can rapidly identify healthy individuals and patients to achieve AD diagnosis. Impurities in clinical samples have little effect on detection. In addition, the ultra-fast detection speed is a key feature of the device. Compared to the classic OECT system, which often requires an incubation time of more than 20 minutes, our system can reach saturation in only two minutes, which is undoubtedly a significant breakthrough for the promotion of OECT technology.

### Monitoring AD progression in an animal model

We further investigated the feasibility of applying the integrated sensor system to monitor the progression of AD in a well-established 5xFAD mouse model. 5xFAD mouse overexpresses human APP and PSEN1 transgenes that include five AD-linked mutations. At 2-4 months of age, these mice rapidly develop amyloid pathology characterized by the accumulation of high levels of intraneuronal Aβ42. As a result, the 5xFAD mice display progressive neuronal loss and cognitive and motor deficiencies ^37-39^. For our experimental workflow (**Fig. 4a**), 5xFAD mice were sacrificed in different age groups (4, 12, and 18 months), with wild-type mice (WT) serving as the control group. After serum collection and perfusion, the whole brain was fixed. The serum samples were diluted and tested using integrated sensor chips modified with anti-Aβ42. High-resolution MRI, considered the gold standard for identifying plague deposition in the brain ^40, 41^, was performed to quantify the age-dependent beta-amyloid loading in each matching brain. Throughout the pathological progression, as shown by the comparison between age groups, there was a significant increase in beta-amyloid loading (MRI results, **Fig. 4b and c**) as well as detected Aβ42+ EV (integrated sensor readouts, **Fig. 4d**). For integrated sensor, additional WT serums collected from age matching pairs were tested. The corresponding *I*_*ds*_ shift was within the healthy baseline level (<0.05), as shown in Supplementary Fig. S5. In the scatter plot representing the pooled data of all 5xFAD mice, we observed a powerful correlation (Pearson *r* = 0.920) between the integrated sensor measurement (normalized I_ds_ shift) and MRI plaque loading (**Fig. 4e**), which suggests that the integrated sensor could be an effective alternative tool of monitoring AD processes. As the role of exosomal Aβ has been extensively studied and explored during the last decade ^28, 42-45^, this proof-of-concept study demonstrated that our integrated OECT sensing system can be applied to enable the diagnosis & prognosis of AD rapidly and minimally invasively.

**Fig. 4.**
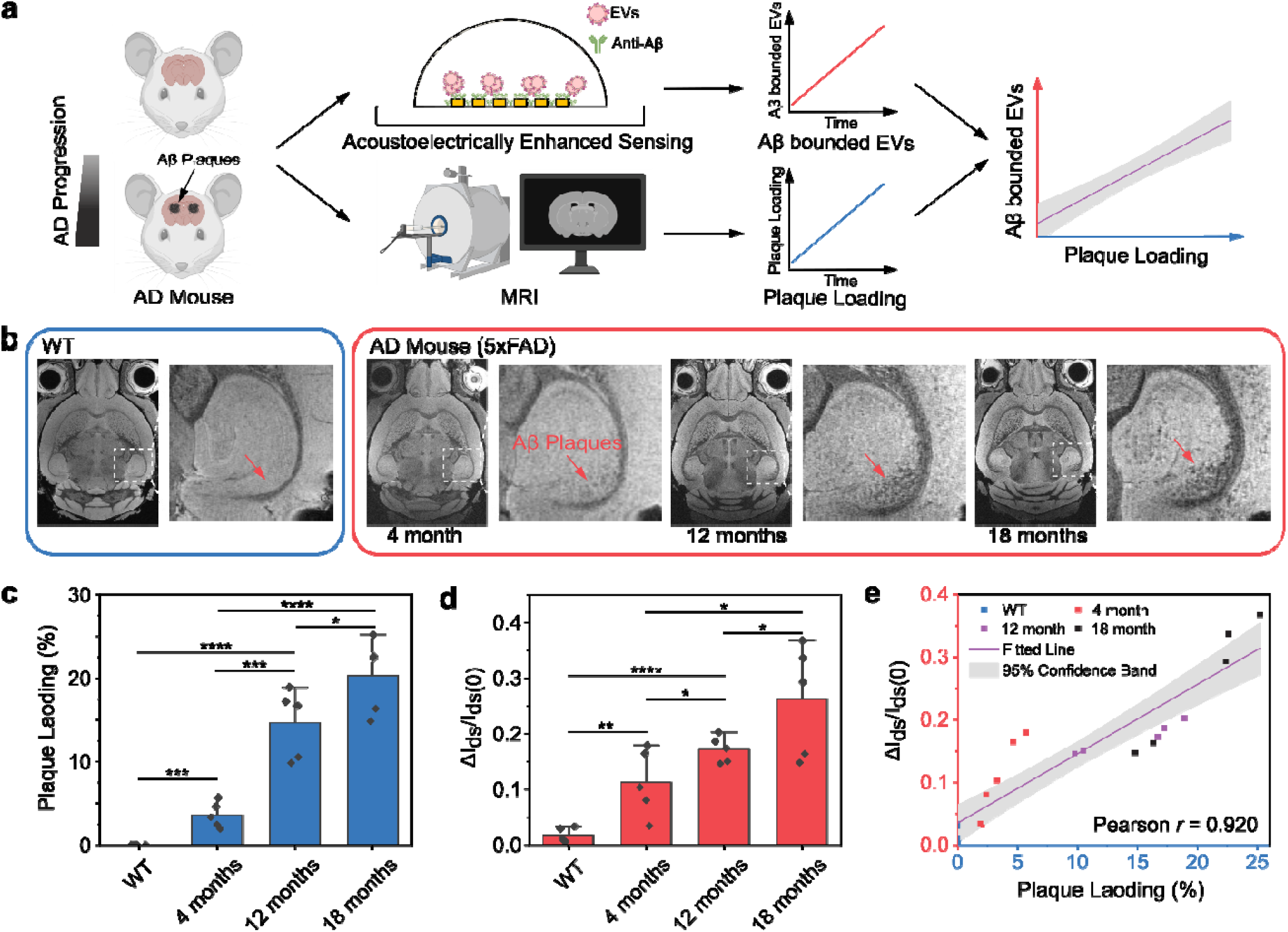
Monitoring of disease progression in AD mouse model. (**a**) The schematic of the monitoring of AD progression in the AD mouse model. The mouse of different ages were examined by MRI and the integrated OECT sensor, respectively, and the profiling of Aβ42+ EV was used to monitor the disease progression in the animal model. **(b)** MRI images of wild type (WT) and AD (5XFAD) mice in 4-18 months. The plaques were gradually increased with age in hippocampus regions (red arrows). **(c)** Quantitative results of plaque loading (%). No plague loading (0%) was detected in WT. **(d)** The results of EV protein expression tested by the integrated OECT sensor in the healthy and AD mouse (**e**) Correlation graph between the integrated OECT sensor and MRI results. A strong correlation was observed from the pooled mice data (20 mice in total). The gray area indicates 95% prediction bands based on linear regression fitting. n=5 independent experiment. Each serum sample was repeated for the EV detection in three independent sensors.

## Discussion

In conclusion, we have developed an ultrafast, sensitive, simple-to-operate portable OECT-based biosensor enhanced by FSAWs and voltage pulse to detect AD-relative EVs. Introducing FSAWs breaks the inefficiency of the classical OECT system in which the target in liquid binds to the antibody on the substrate through Brownian motion, achieving rapid enrichment of EVs. The introduction of the pulsed electric field further accelerates the enrichment process. It improves the performance, shortening the time constant of the OECT-based EV sensing to less than 60 s and decreasing the LOD to hundreds of particles per milliliter. The ultra-fast detection time and high sensitivity allow the integrated OECT sensor to rapidly detect clinical samples in a couple of microliters, which is well-matched to the requirements of POCT. Based on this, this system has quickly detected Tau-EVs and Aβ-EV in serum from AD patient and AD mouse respectively for diagnosis and progress monitoring. This technology is expected to bridge the OECT sensor and POCT for applications in diagnosing and monitoring brain-related diseases. It is worth emphasizing that, as a versatile enrichment technique, the integrated sensor can also be applied to the rapid diagnosis and process monitoring of various other diseases by changing the modified antibodies on the substrate, sensing modules, or the target bioparticles (cells, bacteria, and viruses).

## Materials and methods

### Device fabrication

The electrodes for the acoustofluidic device and OCET were fabricated with standard photolithography by depositing gold (Cr/Au, 10 nm/200 nm) on LiNbO_3_ substrate (500 μm thick, double-side polished, 128° YX-propagation) via a thermal evaporation and liftoff process. As shown in **Fig 1b**, one pair of focused interdigital transducers (IDT) was designed as 16 pairs of finger electrodes with the same finger width and gap (λ/4 = 25 μm) and a 57° focusing angle ^35, 46^. The source-drain of the OECT was located at the focus point patterned as 8 pairs of interdigitated fingers. The length and width of the finger were 325 × 25 μm, and the distance between the organic channel and the side gate electrode was 250 μm. To define the location and dimension of the organic channel, a polyimide masking layer was applied to the electrodes with a window pre-opened with laser cutting. The poly(3,4-ethylenedioxythiophene): poly(styrene sulfonate) (PEDOT: PSS) aqueous solution was doped with divinyl sulfone (5%), (3-glycidyloxypropyl)trimethoxysilane (GOPS, 2%), and ethylene glycol (5%) to improve the conductivity and stability of the organic layer ^47-49^. The mixture was then spin-coated onto the channel area and annealed at 80 °C for 1 hr when flushed with nitrogen. To clean the organic channel before biofunctionalization, the device was washed with 100% EtOH, air-dried, treated with oxygen plasma for 3 min, and incubated with 2% GOPS solution overnight. The device was rewashed with 100% EtOH, dried in an oven at 80 °C for 4 hours, then conjugated with antibodies at 4 °C overnight. 5% BSA was used to block the nonspecific binding sites for at least 30 minutes and kept in the fridge (4 °C) upon usage. Between each step after the antibody conjugation, the device was thoroughly rinsed with 1X PBS three times. A 1.5 mm diameter polycarbonate ring with a thickness of 130 μm was prepared with laser cutting and applied on the OECT area to position the droplet of the sample (2.5 μL)

### System setup

The system consists of the following four modules: the power supply module, acoustic drive module, voltage pulse module, and detection module. Functionally, the power supply module provides electrical power to all modules; the acoustic drive module consists of a signal generator and power amplifier, which is responsible for stimulating the IDT electrode to generate focused acoustic waves; the voltage pulse module is used for generating pulsed voltages to accelerate the binding process; the detection module is a portable electrochemical workstation.

### Donor serum sample preparation

The Indiana Biobank, a part of the Indiana Clinical and Translational Sciences Institute (CTSI), provided whole blood samples from AD patients. Serum was collected by density centrifugation, and the EVs were isolated with a commercially available precipitation kit (ExoQuick, System Biosciences, Palo Alto, CA, US) following the protocol provided by the vendor. The freshly isolated EVs were re-suspended in 1X PBS with different dilution factors for further analysis.

### Device characterization

The acoustofluidic EV focusing was driven by a function generator (TGP3152, Aim TTi, UK) providing a continuous sine RF signal of 37.4 MHz and an amplifier (LZY-22+, Mini-circuit, USA). The acoustofluidic focusing was visualized using fluorescent polystyrene beads (∼ 2 μm diameter, Bangs laboratories Inc, FSDG005 dragon green, Fishers, IN, USA) diluted in 1X PBS and monitored with an inverted microscope (IX-81, Olympus, Japan). The intensity of the focused beads was quantified in ImageJ. The transfer characteristics of the OCET were measured with two source-meter units (Keithley 2612B) with custom control software. The detection area was rinsed with 1X PBS three times between each step. The transfer curve was taken with gate voltage V_G_ = -1V to 1V and V_ds_ = -0.1 V. The relative change of the drain current (I_ds_) was calculated between the enriched detection of EV and the baseline (1X PBS).

### Quantification of the EV size distribution and concentration

The EV samples were further quantified with dynamic light scatter (DLS, Zetasizer Nano-S, Malvern) and nanoparticle tracking analyzer (NTA, Zetaview, Particle Metrix). For the DLS, the diluted samples (100 μL) were loaded into a quartz cuvette (zen 2112) to quantify the intensity particle size distribution (PSD). Each measurement was repeated three times with 12 runs per measurement. The system was pre-calibrated with 100 nm beads for the NTA, and then auto alignment and focus optimization were performed according to the user manual. A 1 ml sample solution was injected into the flow cell, and the size distribution measurement was carried out with the minimum brightness set at 20 and the sensitivity set between 70 and 80.

### Scanning electron microscopy (SEM)

Freshly isolated EVs were vortexed and resuspended in 1X PBS at 5&10^5^ particles/ml. 2.5 μL of the EV solution was loaded onto the device focus area. After the enrichment, the surface of the device was washed with 1X PBS 10 times and fixed in 3% paraformaldehyde solution for 30 min. The samples were then washed with DI water 10 times, serially dehydrated with 30%, 50%, 70%, and 100% ethanol (15 min each), and air-dried for another 30 min under a ventilation hood. To make the sample surface conductive, a 5 nm gold-palladium alloy coating was sputtered with argon as gas for plasma (Denton Desktop V sample preparation system). The SEM images were taken with a field emission scanning electron microscope (FEI Quanta 600).

### Detection process of EVs from serum

The detection process of EVs with target proteins includes the following steps: (1) Serum in microliters collected from the fingertip is diluted at a high level to reduce further the chance of coagulation or non-specific adsorption on the substrate. (2) 5 μL of diluted serum sample is added dropwise to the integrated OECT chip. (3) Turn on the acoustic field and voltage pulse, the enrichment of EVs can be completed within two minutes. (4) Remove the sample and physically absorb biomolecules from the sensing area by rinsing PBS buffer after incubation. (5) Characterize the OECT device performance in the PBS buffer. (6) The data is transferred to the cell phone via Bluetooth. After analysis, a diagnostic report is output on the phone.

### Simulation model

A three-dimensional (3D) numerical model was used to visualize the pattern of SAWs on the surface with COMSOL. 16 pairs of IDTs with a focused angle of 40 degrees were distributed on the surface of the lithium niobate substrate with a width and spacing of 25 μm, as shown in **Fig**.**1c**. In addition to the symmetric interface, the interface was set as a perfect match layer to represent the infinity boundary, and the bottom interface was set as a fixed constraint.

### Animal preparation

Animal experiments were carried out in compliance with the Indiana University Institutional Animal Care and Use Committee. Four wild-type mice at 12 months and five 5xFAD mice at the ages of 4, 12, and 18 months were sacrificed and perfused with the PBS solution. The mouse brains were immersed in buffered formalin for 24 hours and then placed in a PBS solution of 0.5% Prohance (Bracco Diagnostics Inc., Princeton, NJ) to shorten T1 and reduce scan time ^41^.

### MRI protocol

The MRI of the specimens was acquired on a 30-cm bore 9.4 *τ* magnet (Bruker BioSpec 94/30, Billerica, MA) with a maximum gradient strength of 660 mT/m on each axis. A high-sensitivity cryogenic RF surface receive-only coil was used for signal reception (Bruker CryoProbe). A 3D gradient echo (GRE) pulse sequence was performed at the spatial resolution 18 μm^3^ with two TEs (10 and 24.5 ms) ^40^. The matrix size is 900 × 640 × 420. All other parameters were kept the same: FOV = 16.2 × 11.52 × 7.56 mm^3^, flip angle = 45º, bandwidth (BW) = 125 kHz, average=2, and TR = 100 ms. The scan time was about 16 hours.

### Statistical analysis

The statistics comparing different groups were conducted using the t-test. Statistical significance was as follows: *p<0.05, **p<0.01, ***p<0.005. ****p<0.001.

## Acknowledgment

F.G. acknowledges the National Institute of Health Awards (DP2AI160242, R01DK133864, and U01DA056242).

## Supporting Figures and captions

**Supplementary Fig. S1.**
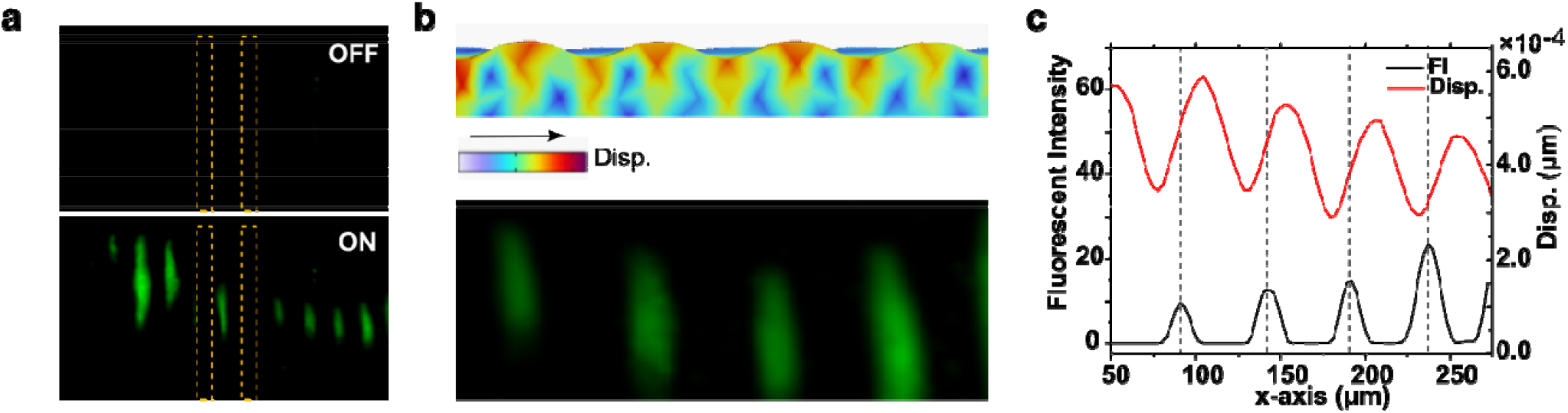
Characterization of acoustic enrichment with 2 μm fluorescent polystyrene microbeads. (a) Distribution of the microbeads with and without acoustic enrichment. (b) Comparison of the acoustic wave simulation results with the actual particle enrichment at the same positions. (c) The curves are extracted from the surface displacements in the simulation and the fluorescence intensity in the fluorescence image at the position of the transverse median.

**Supplementary Fig. S2.**
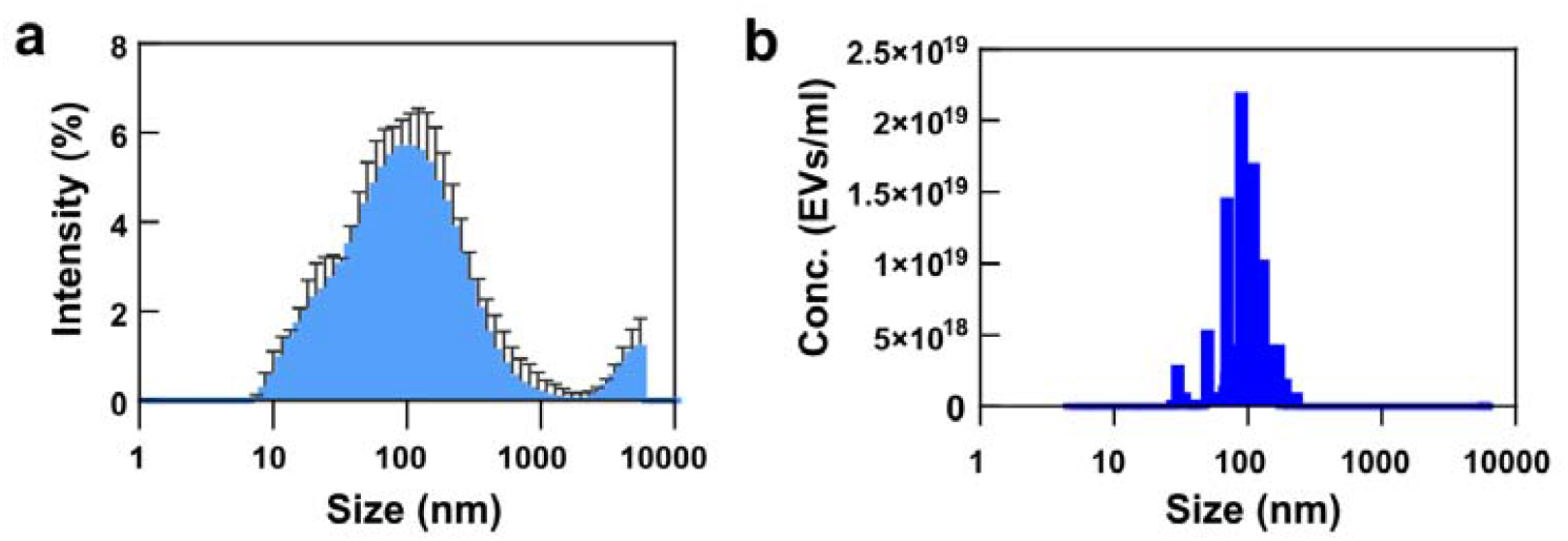
Representative characteristics of purified EVs collected from an AD patient sample. DLS (a) and NTA (b) were used to study the size distribution and total concentration.

**Supplementary Fig. S3.**
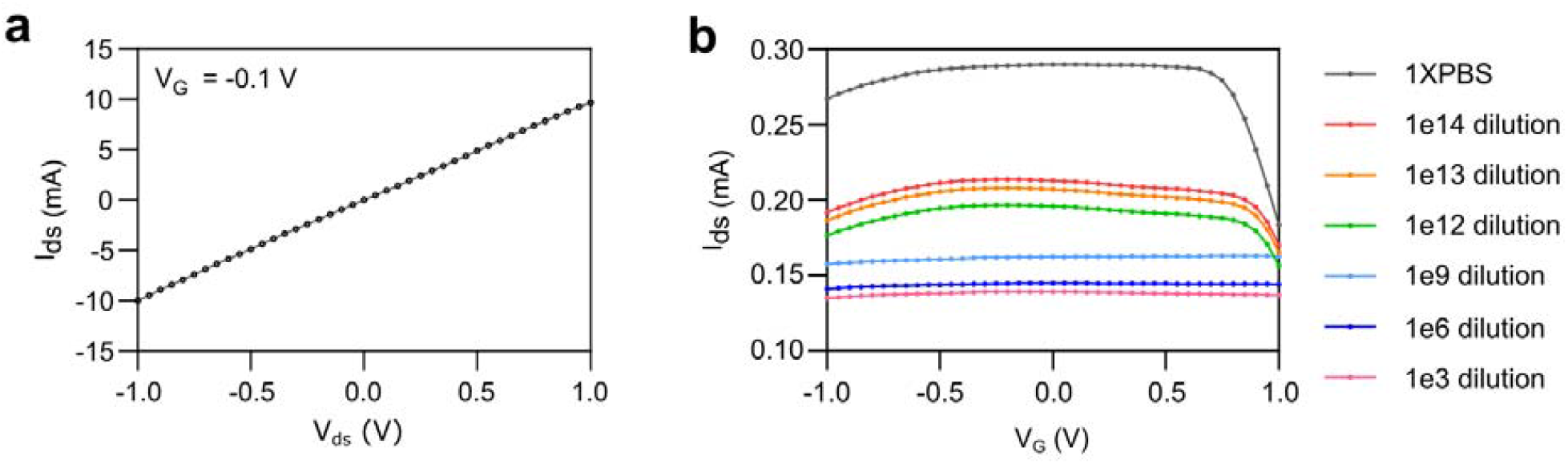
Characteristics of the processed OECT sensor. (a) The transfer curve of the baseline showed excellent linearity. (b) The representative transition curve for EV detection in AD patient sample. Each curve indicated a different dilution factor or baseline (1XPBS). The curve shifted further down based on the EV concentration.

**Supplementary Fig. S4.**
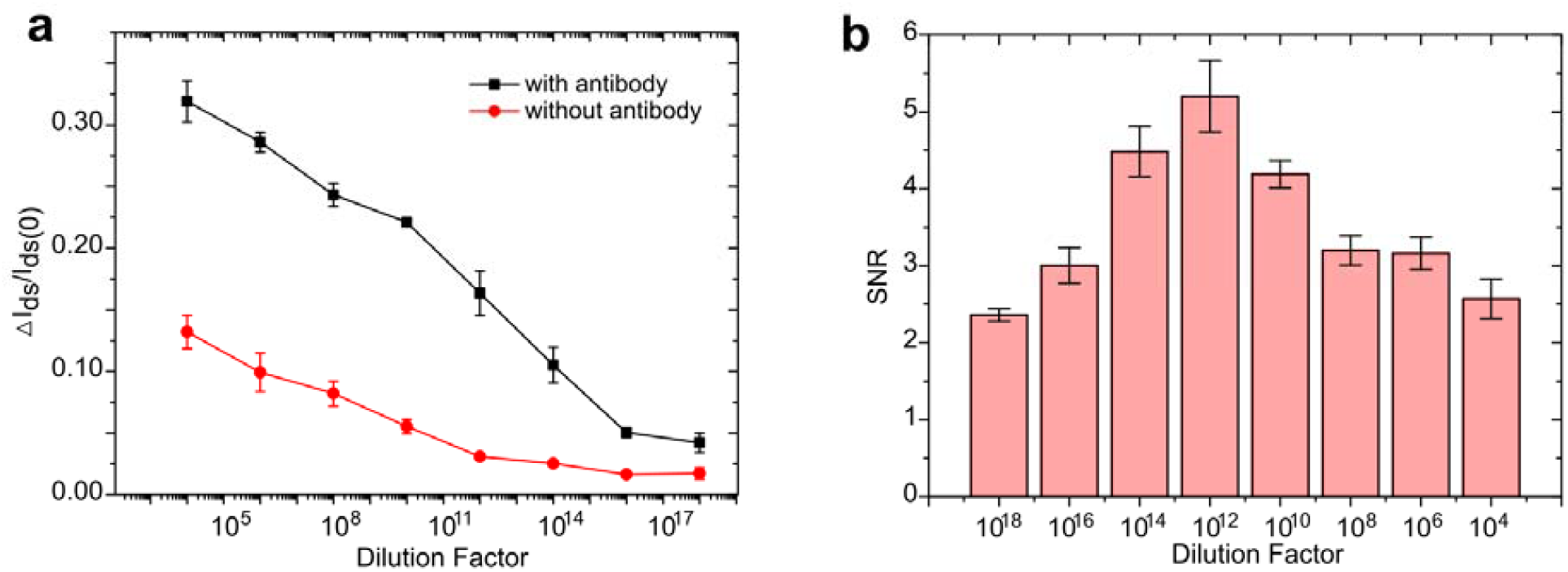
(a) The response of the integrated OECT sensor modified/unmodified antibodies for detecting AD serum in different dilutions. (b) The calculated signal-to-noise ratio (SNR) in various dilutions.

**Supplementary Fig. S5.**
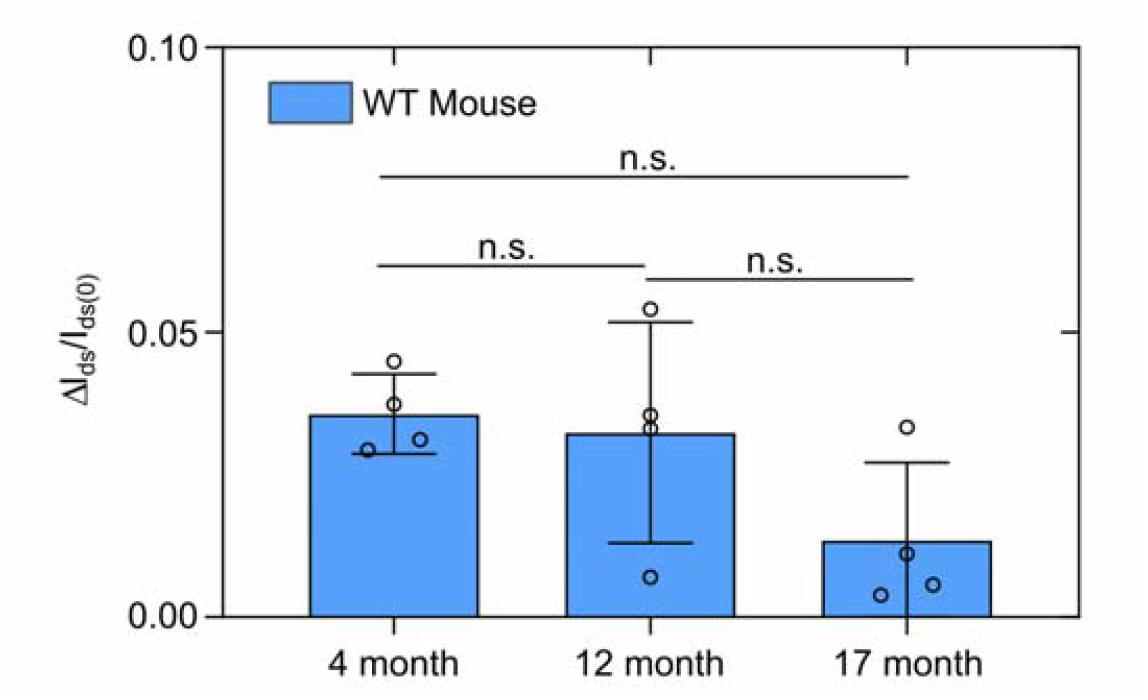
The EV detection of WT aging matching pairs. There was no significant difference between the age groups, and the *I*_*ds*_ shift fell within the healthy baseline range (0.027 ± 0.016).

